# Avoiding potential biases in ses.PD estimations with the Picante software package

**DOI:** 10.1101/579300

**Authors:** Rafael Molina-Venegas

## Abstract

1. Faith’s phylogenetic diversity (PD) is one of the most widespread used indices of phylogenetic structure in the eco-phylogenetic literature. The metric became notably popular with the publication of the function *pd* as part of the Picante R package, which is nowadays a reference software for phylogenetic analyses.
2. Because PD is not statistically independent of species richness, the routine procedure is to standardize the observed PD values for unequal richness across samples. The function *ses.pd*, which is also implemented in the Picante R package, is the reference function to conduct such standardization.
3. Unfortunately, I have detected an anomaly in the function that may result in biased estimations of standardized PD values, particularly in communities with low species richness (i.e. less than four species) and unbalanced phylogenies.
4. I conduct a simple simulation exercise to illustrate the issue and propose two alternative and easy to implement solutions to go around the problem.

## Introduction

Faith’s phylogenetic diversity (PD; Faith, 1992), defined as the sum of all branch lengths connecting taxa in a sample, is one of the most widespread used indices of phylogenetic structure in the eco-phylogenetic literature. The metric became notably popular with the publication of the function *pd* as part of the Picante software (Kembel et al., 2010), which is nowadays a reference R package for phylogenetic analyses. The pd function includes three arguments: (i) “samp”, a community data matrix (samples in rows and species in columns), (ii) “tree”, a phylo tree object including all the species in the community data matrix, and (iii) “include.root”, which is a logical argument. If the latter is set to TRUE (default = TRUE), then the PD of all samples in the community data matrix will include the distance from the most recent common ancestor (MRCA) of the species in each sample and the root of the supplied phylogeny (hereafter “MRCA – root” distance). Otherwise, the MRCA – root distance is excluded from the computations.

Importantly, PD is not statistically independent of species richness, and the routine procedure is to standardize the observed PD values for unequal richness across samples (see documentation for the *pd* function in Picante R package, Kembel et al., 2010). Typically, the observed PD is compared to a null distribution of PD values generated by shuffling species names across the phylogenetic tips a high number of times (e.g. 999 times). The function *ses.pd*, which is also implemented in the Picante software, is the reference function to conduct such standardization. In addition to the arguments of *pd, ses.pd* includes multiple null models that can be used to generate null distributions (the default model is “taxa.labels”, which shuffles taxa labels across the phylogenetic tips). Since *ses.pd* calls internally to *pd*, the user can specify if the MRCA – root distance should be included in the calculation of the observed PD and the corresponding null PD values. However, I have noted that *ses.pd* computes null PD values without including the MRCA – root distance regardless of the logical value that is specified in the include.root argument. This is, if include.root = TRUE (default = TRUE), the observed PD will include the MRCA – root distance, but the null PD values will not (see Supplementary Material). Unfortunately, this anomaly in the *ses.pd* function may result in biased estimations of standardized PD values (hereafter “ses.PD”), particularly in samples with low species richness. This is because the lower the species richness in the samples, the lower the probability for the phylogenetic branches connecting species in the null samples to traverse the root node of the supplied phylogeny, and therefore the higher the impact of excluding the MRCA – root distance from the computations when it should be included (i.e. when include.root = TRUE). Here, I conducted a simple simulation exercise to illustrate this issue.

## Materials and Methods

I simulated four different community data matrices with n = 50 samples (rows) and m = 25, 50, 100 and 200 species (columns) respectively. Then, I used the function *pbtree* implemented in Phytools R package (Revell, 2012) to simulate 500 pure-birth phylogenies (root to tip distance scaled to unit) of m = 25, 50, 100 and 200 tips, respectively (2000 phylogenies in total), representing the species in the community data matrices. Finally, I simulated four different community datasets per community data matrix (each community data matrix represents a different species pool) with fixed species richness within datasets (i.e. equal row sums). To do so, I assigned n = 2, 4, 8 and 16 species, respectively, to each of samples in the community data matrices by randomly picking species from the corresponding pools. For each dataset (16 in total), I computed ses.PD values for the samples using 500 simulated phylogenies and the *ses.pd* function as implement in Picante (Kembel et al., 2010, hereafter “*ses.pd*-Picante”). The argument include.root was set to TRUE, and null distributions were generated using the taxa.labels model and 999 randomizations. Then, I reanalyzed the data using a corrected version of the *ses.pd*-Picante function that actually includes the MRCA – root distance in all the computations if the argument include.root is set to TRUE. Both functions are identical in all other respects (see Supplementary Material). I used the same seed (random number generator) to analyze the data with both functions. Finally, I compared the ses.PD values derived from each function using cross-validation R-squared (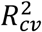) (Molina-Venegas, Moreno-Saiz, Castro, Davies, Peres-Neto & Rodríguez, 2018). 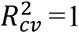 indicates perfect match between ses.PD values obtained from both functions, and 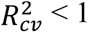 indicates imperfect match. 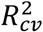 varies from 1 to minus infinity. Since I used the same seed to analyze the data with both functions, the randomization pattern was preserved, and therefore 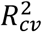 will be equal to 1 in case both functions yield identical results. The R code to reproduce all the analyses along with the corrected version of the *ses.pd*-Picante function is provided as Supplementary Material. All analyses were conducted using Picante version 1.7 (latest version) and R version 3.4.3 (R Core Team., 2017), yet results were the same regardless of the version of the package (the *ses.pd* function was first implemented in Picante version 0.7-2 and delivered in CRAN R repository in July 2009).

## Results and Discussion

I found substantial mismatch between the ses.PD values derived from the *ses.pd*-Picante function and its corrected version, particularly at low species richness and regardless of the size of the species pool (Figs. 1 and S1 in Appendix 1). More specifically, the *ses.pd*-Picante function yielded higher ses.PD values than expected (i.e. above the 1:1 line) in the negative side of the distribution (Figs. 2 and S2 in Appendix 1). However, results derived from both functions rapidly converged following an exponential trend as species richness increased (Figs. 1 and S1 in Appendix 1), suggesting that the *ses.pd*-Picante function will introduce biases only when species richness is very low (i.e. less than four species). Fortunately, such low-richness levels are rare in natural communities, yet they are eventually reported along with their ses.PD values (e.g. Mennes, Moerland, Rath, Smets & Merckx, 2015; Geedicke, Schultz, Rudolph & Oldeland, 2016; Nowakowski, Frishkoff, Thompson, Smith & Todd, 2018). On the other hand, some simulation analyses have also reported ses.PD values for samples including only two species (e.g. Mazel et al., 2016), and diversity experiments often include plots with very few species (e.g. Symstad et al., 2003).

**Figure 1.**
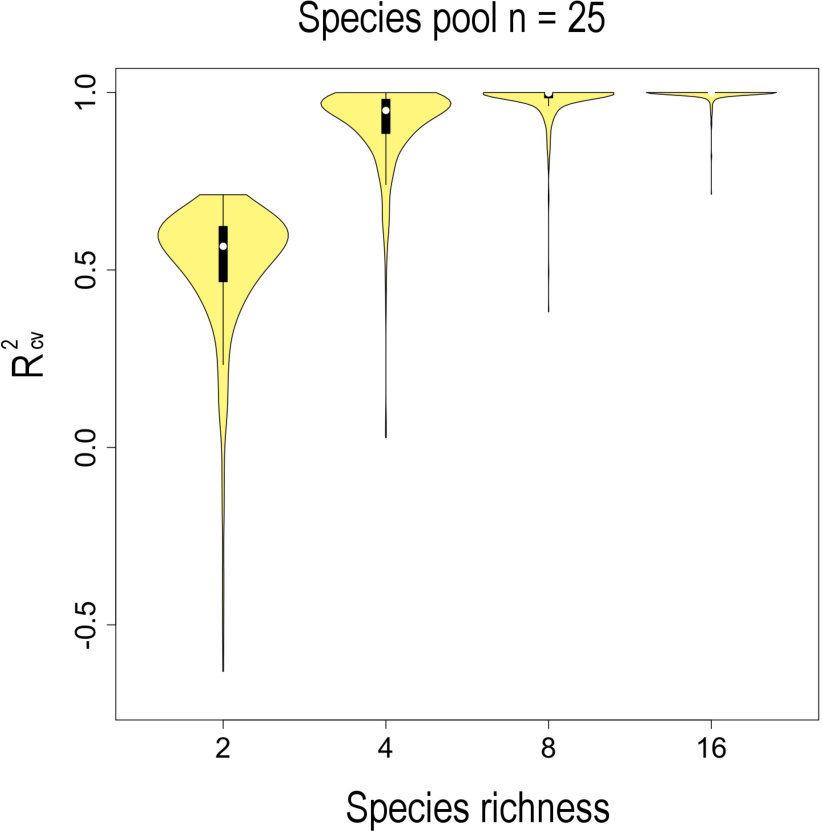
Violin plot showing the cross-validation R-squared scores for the comparisons between ses.PD values derived from the function *ses.pd*-Picante (Kembel et al., 2010) and its corrected version. Analyses were conducted using datasets with species richness = 2, 4, 8 and 16, respectively, 500 simulated phylogenies and a species pool (community data matrix) of n = 25 species (see Fig. S1 in Appendix 1 for results derived from community data matrices with n = 50, 100 and 200 species).

**Figure 2.**
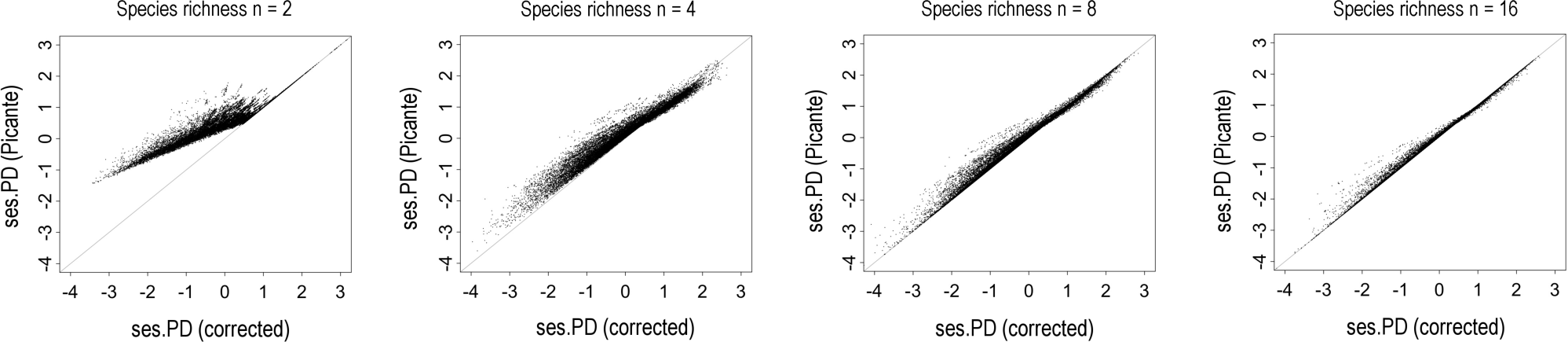
Scatter plots showing the relationship between ses.PD values (25,000 per plot) derived from the function *ses.pd*-Picante (Kembel et al., 2010, y-axis) and its corrected version (x-axis). Analyses were conducted using datasets with species richness n = 2, 4, 8 and 16, respectively, 500 simulated phylogenies and a species pool of 25 species (see Fig. S2 for results derived from species pools of 50, 100 and 200 species). The grey lines represent the expected 1:1 relationship.

Figs. 3 and S3 show that mismatches were more likely to occur with highly unbalanced trees (i.e. those with internal nodes defining divergent lineages of unequal size). This is because the higher the imbalance of the phylogeny, the lower the probability for the phylogenetic branches connecting species in the null samples to traverse the root node of the supplied phylogeny, and therefore the higher the impact of excluding the MRCA – root distance when it should be included. Given the unbalanced nature of most real phylogenies, I conclude that future studies will avoid potential biases in ses.PD estimations (particularly in communities with very low species richness) by either removing the MRCA – root distance from all the computations conducted by the *ses.pd*-Picante function (i.e. setting the include.root argument to FALSE) or using its corrected version if the MRCA – root distance is to be included (see Supplementary Material).

**Figure 3.**
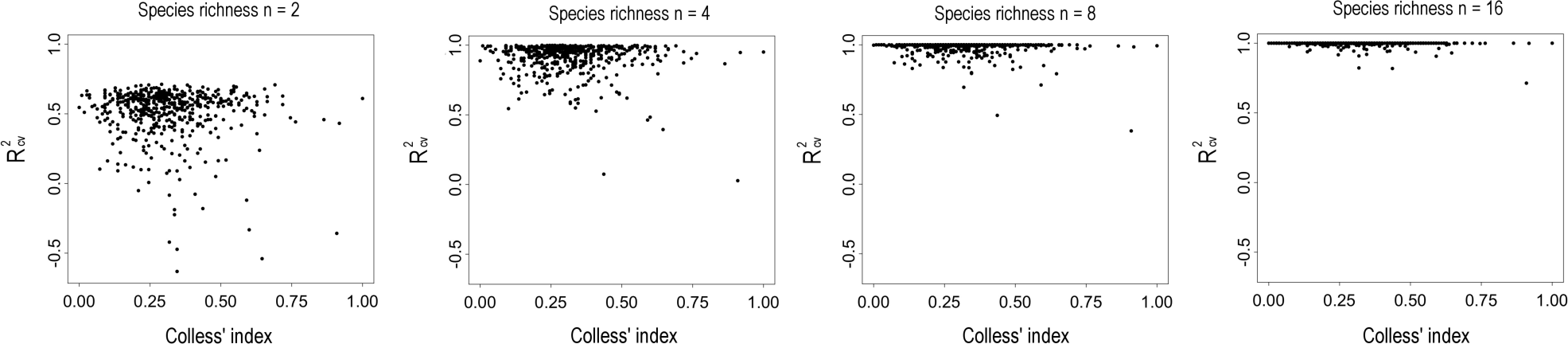
Relationship between cross-validation R-squared scores (y-axis) and tree imbalance (i.e. Colless’ index, x-axis). Analyses were conducted using datasets with species = 2, 4, 8 and 16, respectively, 500 simulated phylogenies and a species pool of 25 species (see Fig. S3 for results derived from species pools of 50, 100 and 200 species). The higher the value of the Colless’ index, the higher the imbalance of the phylogeny (values scaled between 0 and 1).

## Acknowledgments

I thank the Scientific Computation Centre of Andalusia (CICA) for the computing services they provided.

## Supporting Information

**Appendix 1.** R code used for the analyses.

## REFERENCES

Faith, D. P. (1992). Conservation evaluation and phylogenetic diversity. Biological Conservation, 61, 1–10. doi:10.1016/0006-3207(92)91201-3

Geedicke, I., Schultz, M., Rudolph, B., & Oldeland, J. (2016). Phylogenetic clustering found in lichen but not in plant communities in European heathlands. Community Ecology, 17, 216–224. doi:10.1556/168.2016.17.2.10

Kembel, S.W., Cowan, P. D., Helmus, M. R., Cornwell, W. K., Morlon, H, Ackerly, D. D., …, Webb, C. O. (2010). Picante: R tools for integrating phylogenies and ecology. Bioinformatics, 26, 1463–1464.

Mazel, F., Davies, T. J., Gallien, L., Renaud, J., Groussin, M., Münkemüller, T., & Thuiller, W. (2016). Influence of tree shape and evolutionary time-scale on phylogenetic diversity metrics. Ecography, 39, 913–920. doi:10.1111/ecog.01694

Mennes, C. B., Moerland, M. S., Rath, M., Smets, E. F., & Merckx, V. S. F. T. (2015). Evolution of mycoheterotrophy in Polygalaceae: The case of Epirixanthes. American Journal of Botany, 102, 598–608. doi:10.3732/ajb.1400549

Molina-Venegas, R., Moreno-Saiz, J. C., Parga, I. C., Davies, T. J., Peres-Neto, P. R., & Rodríguez, M. Á. (2018). Assessing among-lineage variability in phylogenetic imputation of functional trait datasets. Ecography, 41, 1740–1749 doi:10.1111/ecog.03480

Nowakowski, A. J., Frishkoff, L. O., Thompson, M. E., Smith, T. M., & Todd, B. D. (2018). Phylogenetic homogenization of amphibian assemblages in human-altered habitats across the globe. Proceedings of the National Academy of Sciences, 115, E3454–E3462. doi:10.1073/pnas.1714891115

R Core Team (2017) R: a language and environment for statistical computing. R Foundation for Statistical Computing, Vienna, Austria.

Revell, L. J. (2012). phytools: an R package for phylogenetic comparative biology (and other things). Methods in Ecology and Evolution, 3, 217–223.

Symstad, A. J., Chapin, F. S., Wall, D. H., Gross, K. L., Huenneke, L. F., Mittelbach, G. G., … Tilman, D. (2003). Long-term and large-scale perspectives on the relationship between biodiversity and ecosystem functioning. BioScience, 53, 89–98. doi:10.1641/0006-3568(2003)053[0089:LTALSP]2.0.CO;2

