## Appendix 1 for "Avoiding potential biases in ses.PD estimations with the Picante software package"

### The "include.root" argument of the *ses.pd* function, a pocket example

In this section, I make use of the example dataset included in the Picante R package to show the anomaly of the *ses.pd* function (Kembel et al., 2010, hereafter "*ses.pd*-Picante") described in this paper. For this purpose, I used the Picante version 1.7 (latest version), yet results were the same regardless of the version of the package. The *ses.pd*-Picante function was first implemented in version 0.7-2 and delivered in 2009.

### load the package

```
require(picante)
```

### load the example dataset

```
data(phylocom)
```

### extract the community data matrix

```
phylocom$sample -> Sample
```

### extract the phylogeny

```
phylocom$phylo -> Tree
```

### prune the phylogeny to retain only the species in the community data matrix

```
prune.sample(Sample,Tree)->Tree_F
```

### sort species names in the community data matrix in the same order as the phylogenetic tips

```
Sample <- Sample[,Tree_F$tip.label]
```

### fix the seed

```
set.seed(12345)
```

### compute ses.PD values excluding the MRCA-root distance with the *ses.pd*-Picante function implemented in Picante (analysis 1)

```
ses.pd(Sample,Tree_F,null.model="taxa.labels",include.root=FALSE,runs=999) ->
Picante_no_root_1
```

### fix the seed

```
set.seed(12345)
```

### compute ses.PD values including the MRCA-root distance with the *ses.pd*-Picante function implemented in Picante (analysis 2)

```
ses.pd(Sample,Tree_F,null.model="taxa.labels",include.root=TRUE,runs=999)->
Picante_root_2
```

**Table 1.** Results excluding the MRCA-root distance with the *ses.pd*-Picante function implemented in Picante (analysis 1).

| sample | ntaxa | pd.obs | pd.rand.mean | pd.rand.sd | ses.PD | runs |
|---|---|---|---|---|---|---|
| clump1 | 8 | 14 | 26,08308308 | 1,720472073 | -7,0231207 | 999 |
| clump2a | 8 | 16 | 26,07107107 | 1,777152356 | -5,6669711 | 999 |
| clump2b | 8 | 18 | 26,20020020 | 1,646019051 | -4,9818380 | 999 |
| clump4 | 8 | 22 | 26,05805806 | 1,779597304 | -2,2803238 | 999 |
| even | 8 | 30 | 26,11411411 | 1,767835251 | 2,19810408 | 999 |
| random | 8 | 27 | 26,12612613 | 1,768151812 | 0,49423012 | 999 |

**Table 2.** Results including the MRCA-root distance with the *ses.pd*-Picante function implemented in Picante (analysis 2).

| sample | ntaxa | pd.obs | pd.rand.mean | pd.rand.sd | ses.PD | runs |
|---|---|---|---|---|---|---|
| clump1 | 8 | 16 | 26,08308308 | 1,720472073 | -5,8606491 | 999 |
| clump2a | 8 | 17 | 26,07107107 | 1,777152356 | -5,1042732 | 999 |
| clump2b | 8 | 18 | 26,20020020 | 1,646019051 | -4,9818380 | 999 |
| clump4 | 8 | 22 | 26,05805806 | 1,779597304 | -2,2803238 | 999 |
| even | 8 | 30 | 26,11411411 | 1,767835251 | 2,19810408 | 999 |
| random | 8 | 27 | 26,12612613 | 1,768151812 | 0,49423012 | 999 |

Note that regardless of the logical value indicated for the include.root argument, the descriptive statistics for the null distributions (i.e. mean and standard deviation) are identical, but they should not be. In contrast, the observed PD for the samples "clump1" and "clump2a" differ between both analyses (in red), because in the second analysis (Table 2) the MRCA-root distance is included in the calculations. The samples "clumps2b", "clump4", "even" and "random" do not show differences in their observed PD values between both analyses because the phylogenetic branches connecting their constituent species traverse the root node of the supplied phylogeny (see Figure A1)

**Figure A1.** Arrangement of the species in the six community samples analyzed (example dataset in Picante) across the tips of the supplied phylogeny. The phylogenetic branches connecting the species in each community are in blue (the sum up of these branches is the observed PD of the samples excluding the MRCA-root distance). Note that the phylogenetic branches connecting species in samples “clump2b”, “clump4”, “even” and “random” traverse the root node of the supplied phylogeny, and therefore their MRCA-root distances (in orange) are equal to zero. The vertical grey bar represents 1 unit of PD.

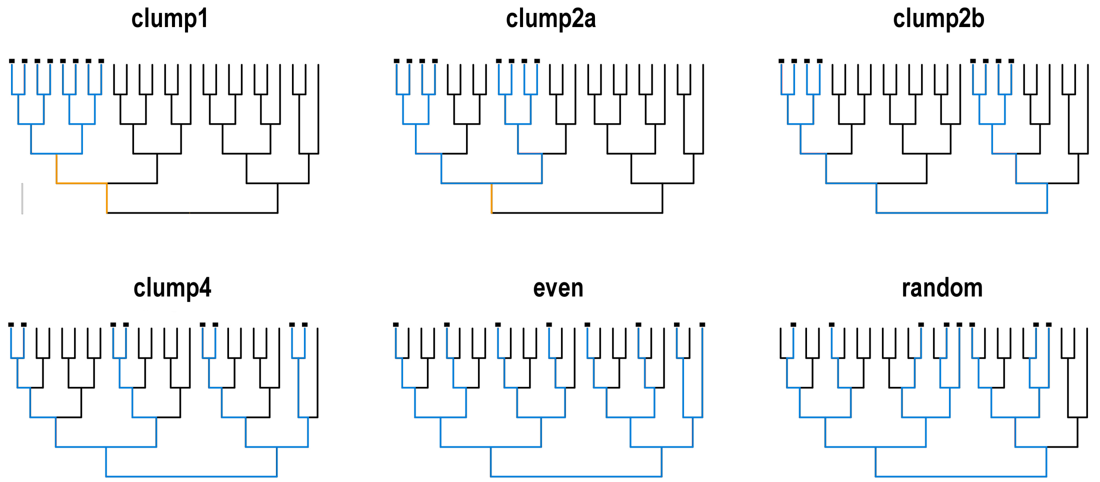


Now, I use the corrected version of the *ses.pd*-Picante function that actually includes the MRCA – root distance in all the computations as long as the argument include.root is set to TRUE.

### source to the corrected version of the *ses.pd*-Picante function (must be located in the current working directory)

```
source("ses_pd_fixed.txt")
```

### fix the seed

```
set.seed(12345)
```

### compute ses.PD values excluding the MRCA-root distance with the corrected version of the *ses.pd*-Picante function (analysis 3)

```
ses.pd.fixed(Sample,Tree_F,null.model="taxa.labels",include.root=FALSE,runs=99
9) -> Picante_no_root_3
```

### fix the seed

```
set.seed(12345)
```

### compute ses.PD values including the MRCA-root distance with the corrected version of the *ses.pd*-Picante function (analysis 4)

```
ses.pd.fixed(Sample,Tree_F,null.model="taxa.labels",include.root=TRUE,runs=999
)-> Picante_root_4
```

**Table 3.** Results excluding the MRCA-root distance with the corrected version of the *ses.pd*-Picante function (analysis 3)

| sample | ntaxa | pd.obs | pd.rand.mean | pd.rand.sd | ses.pd | runs |
|---|---|---|---|---|---|---|
| clump1 | 8 | 14 | 26,08308308 | 1,720472073 | -7,0231207 | 999 |
| clump2a | 8 | 16 | 26,07107107 | 1,777152356 | -5,6669711 | 999 |
| clump2b | 8 | 18 | 26,20020020 | 1,646019051 | -4,9818380 | 999 |
| clump4 | 8 | 22 | 26,05805806 | 1,779597304 | -2,2803238 | 999 |
| even | 8 | 30 | 26,11411411 | 1,767835251 | 2,19810408 | 999 |
| random | 8 | 27 | 26,12612613 | 1,768151812 | 0,49423012 | 999 |

**Table 4.** Results including the MRCA-root distance with the corrected version of the *ses.pd*-Picante function (analysis 4)

| sample | ntaxa | pd.obs | pd.rand.mean | pd.rand.sd | ses.pd | runs |
|---|---|---|---|---|---|---|
| clump1 | 8 | 16 | 26,08808808 | 1,703541122 | -5,9218342 | 999 |
| clump2a | 8 | 17 | 26,07507507 | 1,762834235 | -5,1480025 | 999 |
| clump2b | 8 | 18 | 26,20320320 | 1,636185796 | -5,0136135 | 999 |
| clump4 | 8 | 22 | 26,06406406 | 1,760710112 | -2,3081960 | 999 |
| even | 8 | 30 | 26,11811811 | 1,753342670 | 2,21398929 | 999 |
| random | 8 | 27 | 26,13413413 | 1,743590043 | 0,49659946 | 999 |

Now, the descriptive statistics for the null distributions (i.e. mean and standard deviation, in red) between analyses 3 and 4 are not identical, because the corrected version of the *ses.pd*-Picante function actually includes the MRCA-root distance in all the computations (i.e. to compute either observed PD and null PD values) in analysis 4, where the argument include.root was set to TRUE. Note that results from analyses 1 and 3 (include.root = FALSE) are identical. In this example, the impact of the anomaly detected for the *ses.pd*-Picante function is rather negligible, since species richness is equal to 8 in all cases. Finally, we can add two extra samples to the dataset with n = 2 and 3 closely-related species, respectively, in order to illustrate the potential impact of the anomaly.

### create the new samples including n = 2 and 3 species, respectively

```
SR2 <- c(1,1,rep(0,23))

SR3 <- c(1,1,1,rep(0,22))
```

### add the new samples to the dataset

```
Sample2 <- rbind (Sample,SR2,SR3)
```

### fix the seed

```
set.seed(12345)
```

### compute ses.PD values including the MRCA-root distance with the *ses.pd*-Picante function implemented in Picante (analysis 5)

```
ses.pd(Sample2,Tree_F,null.model="taxa.labels",include.root=TRUE,runs=999)->
Picante_root_5
```

### fix the seed

```
set.seed(12345)
```

### compute ses.PD values including the MRCA-root distance with the corrected version of the *ses.pd*-Picante function (analysis 6)

```
ses.pd.fixed(Sample2,Tree_F,null.model="taxa.labels",include.root=TRUE,runs=99
9)-> Picante_root_6
```

### fix the seed

```
set.seed(12345)
```

### compute ses.PD values excluding the MRCA-root distance with the *ses.pd*-Picante function implemented in Picante (analysis 7)

```
ses.pd(Sample2,Tree_F,null.model="taxa.labels",include.root=FALSE,runs=999) ->
Picante_no_root_7
```

### fix the seed

```
set.seed(12345)
```

### compute ses.PD values excluding the MRCA-root distance with the corrected version of the *ses.pd*-Picante function (analysis 8)

```
ses.pd.fixed(Sample2,Tree_F,null.model="taxa.labels",include.root=FALSE,runs=9
99) -> Picante_no_root_8
```

**Table 5.** Results including the "MRCA-root" distance with the *ses.pd*-Picante function implemented in Picante (analysis 5)

| sample | ntaxa | pd.obs | pd.rand.mean | pd.rand.sd | ses.PD | runs |
|---|---|---|---|---|---|---|
| clump1 | 8 | 16 | 26,0830831 | 1,72047207 | −5,8606491 | 999 |
| clump2a | 8 | 17 | 26,0710711 | 1,77715236 | −5,1042732 | 999 |
| clump2b | 8 | 18 | 26,2002002 | 1,64601905 | −4,981838 | 999 |
| clump4 | 8 | 22 | 26,0580581 | 1,7795973 | −2,2803238 | 999 |
| even | 8 | 30 | 26,1141141 | 1,76783525 | 2,19810408 | 999 |
| random | 8 | 27 | 26,1261261 | 1,76815181 | 0,49423012 | 999 |
| SR2 | 2 | 6 | 8,17217217 | 2,22987243 | −0,974124 | 999 |
| SR3 | 3 | 8 | 12,3363363 | 1,89741703 | −2,2853892 | 999 |

**Table 6.** Results including the MRCA-root distance with the corrected version of the *ses.pd*-Picante function (analysis 6)

| sample | ntaxa | pd.obs | pd.rand.mean | pd.rand.sd | ses.PD | runs |
|---|---|---|---|---|---|---|
| clump1 | 8 | 16 | 26,0880881 | 1,70354112 | −5,9218342 | 999 |
| clump2a | 8 | 17 | 26,0750751 | 1,76283423 | −5,1480025 | 999 |
| clump2b | 8 | 18 | 26,2032032 | 1,6361858 | −5,0136135 | 999 |
| clump4 | 8 | 22 | 26,0640641 | 1,76071011 | −2,308196 | 999 |
| even | 8 | 30 | 26,1181181 | 1,75334266 | 2,2139893 | 999 |
| random | 8 | 27 | 26,1341341 | 1,74359004 | 0,49659946 | 999 |
| SR2 | 2 | 6 | 9,08608609 | 1,11493621 | −2,7679486 | 999 |
| SR3 | 3 | 8 | 12,6486486 | 1,39051888 | −3,3431036 | 999 |

Importantly, the *ses.pd*-Picante function yielded ses.PD values for samples SR2 and SR3 equal to -0.97 and -2.28, respectively (Table 5, in red). However, these values are higher than expected (i.e. above the 1:1 line), because the MRCA-root distance is not included in the computation of null PD values. The actual values are lower (Table 6, in red), and particularly for sample SR2. When the MRCA-root distance is excluded from the computations (analyses 7 and 8), results derived from the *ses.pd*-Picante function and its corrected version are identical (Tables 7 and 8):

**Table 7.** Results excluding the MRCA-root distance with the *ses.pd*-Picante function implemented in Picante (analysis 7)

| sample | ntaxa | pd.obs | pd.rand.mean | pd.rand.sd | ses.PD | runs |
|---|---|---|---|---|---|---|
| clump1 | 8 | 14 | 26,0830831 | 1,72047207 | -7,0231207 | 999 |
| clump2a | 8 | 16 | 26,0710711 | 1,77715236 | -5,6669711 | 999 |
| clump2b | 8 | 18 | 26,2002002 | 1,64601905 | -4,981838 | 999 |
| clump4 | 8 | 22 | 26,0580581 | 1,7795973 | -2,2803238 | 999 |
| even | 8 | 30 | 26,1141141 | 1,76783525 | 2,19810408 | 999 |
| random | 8 | 27 | 26,1261261 | 1,76815181 | 0,49423012 | 999 |
| SR2 | 2 | 2 | 8,17217217 | 2,22987243 | -2,7679486 | 999 |
| SR3 | 3 | 5 | 12,3363363 | 1,89741703 | -3,866486 | 999 |

**Table 8.** Results excluding the MRCA-root distance with the corrected version of the *ses.pd*-Picante function (analysis 8)

| sample | ntaxa | pd.obs | pd.rand.mean | pd.rand.sd | ses.PD | runs |
|---|---|---|---|---|---|---|
| clump1 | 8 | 14 | 26,0830831 | 1,72047207 | -7,0231207 | 999 |
| clump2a | 8 | 16 | 26,0710711 | 1,77715236 | -5,6669711 | 999 |
| clump2b | 8 | 18 | 26,2002002 | 1,64601905 | -4,981838 | 999 |
| clump4 | 8 | 22 | 26,0580581 | 1,7795973 | -2,2803238 | 999 |
| even | 8 | 30 | 26,1141141 | 1,76783525 | 2,19810408 | 999 |
| random | 8 | 27 | 26,1261261 | 1,76815181 | 0,49423012 | 999 |
| SR2 | 2 | 2 | 8,17217217 | 2,22987243 | -2,7679486 | 999 |
| SR3 | 3 | 5 | 12,3363363 | 1,89741703 | -3,866486 | 999 |

**Figure A2.** Arrangement of the species in the communities SR2 and SR3 across the tips of the supplied phylogeny. The phylogenetic branches connecting the species in each community are in blue (the sum up of these branches is the observed PD of the samples excluding the MRCA-root distance, here in orange). The vertical grey bar represents 1 unit of PD.

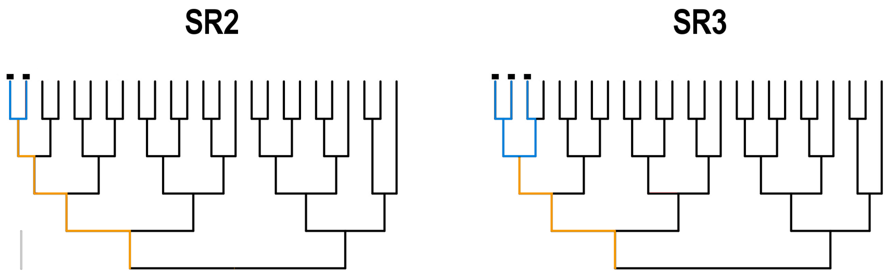

**<u>R code to reproduce all the analyses conducted in this paper</u>**

### Dataset simulation #

```
set.seed(NULL)
Data <- vector(mode="list",length=4)      # dataset store
n_sp = 25       # number of species in the community data matrix (species pool). In the analyses of the paper, this parameter was set to 25, 50, 100 and 200, respectively.
sps <- print(paste("t",1:n_sp,sep="")) # species names
```

### To avoid duplicate rows in the community data matrix (i.e. identical communities), I followed a three-steps procedure:

### 1. Simulate a dataset with n = 100 samples (rows) and m = n_sp species (columns)
### 2. Assign n = Sp_per_comu species to each sample at random from the available in the pool
### 3. Remove duplicate rows and check if there are at least n = 50 non-duplicated samples in the dataset. If this condition is met, the first 50 samples are selected and the resultant dataset is stored. Otherwise, the loop goes back to step 1.

```
Sp_per_comu = 2       # number of species in the samples
Sites=100

for(i in 1:100000){
KK <- matrix(0,nrow=Sites,ncol=n_sp)
colnames(KK)<-sps
row.names(KK)<-paste("site",1:Sites,sep="")
KK <- as.data.frame(KK)
for(u in 1:Sites) {
sample(sps,Sp_per_comu)->Picked
KK[u,colnames(KK)[colnames(KK)%in%Picked]]<-1
}
KK[!duplicated(KK), ]->Samp_F      # remove duplicate samples
if(dim(Samp_F)[[1]]>49){           # are there at least 50 non-duplicated samples?
Samp_F[c(1:50),]->Samp_F
paste("site",1:50,sep="")->row.names(Samp_F)
Samp_F->Data[[1]]
break} else           # loop i is broken when the condition is met and the dataset is stored in the Data object
{1}
}
```

### Same procedure with 4 species per sample

```
Sp_per_comu = 4
for(i in 1:100000){
KK <- matrix(0,nrow=Sites,ncol=n_sp)
colnames(KK)<-sps
row.names(KK)<-paste("site",1:Sites,sep="")
KK <- as.data.frame(KK)
for(u in 1:Sites) {
sample(sps,Sp_per_comu)->Picked
KK[u,colnames(KK)[colnames(KK)%in%Picked]]<-1
}
KK[!duplicated(KK), ]->Samp_F
if(dim(Samp_F)[[1]]>49){
Samp_F[c(1:50),]->Samp_F
paste("site",1:50,sep="")->row.names(Samp_F)
Samp_F->Data[[2]]
break} else
{1}
}
```

### Same procedure with 8 species per sample

```
Sp_per_comu = 8
for(i in 1:100000){
KK <- matrix(0,nrow=Sites,ncol=n_sp)
colnames(KK)<-sps
row.names(KK)<-paste("site",1:Sites,sep="")
KK <- as.data.frame(KK)
for(u in 1:Sites) {
sample(sps,Sp_per_comu)->Picked
KK[u,colnames(KK)[colnames(KK)%in%Picked]]<-1
}
KK[!duplicated(KK), ]->Samp_F
if(dim(Samp_F)[[1]]>49){
Samp_F[c(1:50),]->Samp_F
paste("site",1:50,sep="")->row.names(Samp_F)
Samp_F->Data[[3]]
break} else
{1}
}
```

### Same procedure with 16 species per sample

```
Sp_per_comu = 16
for(i in 1:100000){
KK <- matrix(0,nrow=Sites,ncol=n_sp)
```

```
colnames(KK)<-sps
row.names(KK)<-paste("site",1:Sites,sep="")
KK <- as.data.frame(KK)
for(u in 1:Sites) {
sample(sps,Sp_per_comu)->Picked
KK[u,colnames(KK)[colnames(KK)%in%Picked]]<-1
}
KK[!duplicated(KK), ]->Samp_F
if(dim(Samp_F)[[1]]>49){
Samp_F[c(1:50),]->Samp_F
paste("site",1:50,sep="")->row.names(Samp_F)
Samp_F->Data[[4]]
break} else
{1}
}
```

### Phylogeny simulation #

```
require(phytools)
Pool = 500
```

### number of phylogenies to be simulated. **NOTE**: analyzing the data with the default parameters in the present code will take a very long time. You may consider a lower number of trees (e.g. 50 instead of 500) and/or a lower number of randomizations (e.g. 99 instead of 999) to speed up the analyses.

```
Tree_List <- vector(mode="list",length=Pool)      # Trees store
Size=n_sp      # number of species in the community data matrix (species pool)
for(i in 1:Pool){
pbtree(n=Size,scale=1)->Tree.i
Tree_List[[i]]<-Tree.i
print(paste("Tree ",i," completed",sep=""))
}
```

### ses.PD analyses #

### Analyses with the *ses.pd*-Picante function (Kembel et al. 2010)

```
require(picante)
set.seed(12345) # fix the seed

Data[[1]]->Sample              # dataset with species richness (in the samples) equal to 2
Results_Picante_2sp <- vector(mode="list",length=Pool)  # results store (SR = 2)
for(i in 1:Pool){
Tree_List[[i]]->Tree.i
```

```
Sample.i <- Sample[,Tree.i$tip.label]
ses.pd(Sample.i,Tree.i, include.root=TRUE,runs=999)->Results_Picante_2sp[[i]]
}

Data[[2]]->Sample              # dataset with species richness (in the samples) equal to 4
Results_Picante_4sp <- vector(mode="list",length=Pool)  # results store (SR = 4)
for(i in 1:Pool){
Tree_List[[i]]->Tree.i
Sample.i <- Sample[,Tree.i$tip.label]
ses.pd(Sample.i,Tree.i, include.root=TRUE,runs=999)->Results_Picante_4sp[[i]]
}

Data[[3]]->Sample              # dataset with species richness (in the samples) equal to 8
Results_Picante_8sp <- vector(mode="list",length=Pool)  # results store (SR = 8)
for(i in 1:Pool){
Tree_List[[i]]->Tree.i
Sample.i <- Sample[,Tree.i$tip.label]
ses.pd(Sample.i,Tree.i, include.root=TRUE,runs=999)->Results_Picante_8sp[[i]]
}

Data[[4]]->Sample               # dataset with species richness (in the samples) equal to 16
Results_Picante_16sp <- vector(mode="list",length=Pool)  # results store (SR = 16)
for(i in 1:Pool){
Tree_List[[i]]->Tree.i
Sample.i <- Sample[,Tree.i$tip.label]
ses.pd(Sample.i,Tree.i, include.root=TRUE,runs=999)->Results_Picante_16sp[[i]]
}
```

### Re-analyzing the data the with the corrected version of the *ses.pd*-Picante function

```
set.seed(12345)                # fix the seed (same seed used earlier)
source("ses_pd_fixed.txt")  # source to the ses.pd-fixed function (must be located in the
current working directory)

Data[[1]]->Sample              # dataset with species richness (in the samples) equal to 2
Results_Fixed_2sp <- vector(mode="list",length=Pool)      # results store (SR = 2)

for(i in 1:Pool){
Tree_List[[i]]->Tree.i
Sample.i <- Sample[,Tree.i$tip.label]
ses.pd.fixed(Sample.i,Tree.i, null.model = "taxa.labels",
include.root=TRUE,runs=999)->Results_Fixed_2sp[[i]]
```

```
}

Data[[2]]->Sample            # dataset with species richness (in the samples) equal to 4
Results_Fixed_4sp <- vector(mode="list",length=Pool)  # results store (SR = 4)
for(i in 1:Pool){
Tree_List[[i]]->Tree.i
Sample.i <- Sample[,Tree.i$tip.label]
ses.pd.fixed(Sample.i,Tree.i, null.model = "taxa.labels",
include.root=TRUE,runs=999)->Results_Fixed_4sp[[i]]
}

Data[[3]]->Sample            # dataset with species richness (in the samples) equal to 8
Results_Fixed_8sp <- vector(mode="list",length=Pool)  # results store (SR = 8)
for(i in 1:Pool){
Tree_List[[i]]->Tree.i
Sample.i <- Sample[,Tree.i$tip.label]
ses.pd.fixed(Sample.i,Tree.i, null.model = "taxa.labels",
include.root=TRUE,runs=999)->Results_Fixed_8sp[[i]]
}

Data[[4]]->Sample            # dataset with species richness (in the samples) equal to 16
Results_Fixed_16sp <- vector(mode="list",length=Pool)  # results store (SR = 16)
for(i in 1:Pool){
Tree_List[[i]]->Tree.i
Sample.i <- Sample[,Tree.i$tip.label]
ses.pd.fixed(Sample.i,Tree.i, null.model = "taxa.labels",
include.root=TRUE,runs=999)->Results_Fixed_16sp[[i]]
}
```

### Calculation of cross-validation R-squared scores #

```
R2_2sp<- vector(mode="numeric",length=Pool)     # R^2 cv store (SR = 2)
for(i in 1:Pool){
1-((sum((Results_Fixed_2sp[[i]][,6] -
Results_Picante_2sp[[i]][,6])^2))/length(Results_Fixed_2sp[[i]][,6]))/var(Resu
lts_Fixed_2sp[[i]][,6])->R2_2sp[i]
}

R2_4sp<- vector(mode="numeric",length=Pool)     # R^2 cv store (SR = 4)
for(i in 1:Pool){
1-((sum((Results_Fixed_4sp[[i]][,6] -
Results_Picante_4sp[[i]][,6])^2))/length(Results_Fixed_4sp[[i]][,6]))/var(Resu
lts_Fixed_4sp[[i]][,6])->R2_4sp[i]
```

```
}

R2_8sp<- vector(mode="numeric",length=Pool)   # R2 cv store (SR = 8)
for(i in 1:Pool){
1-((sum((Results_Fixed_8sp[[i]][,6] -
Results_Picante_8sp[[i]][,6])^2))/length(Results_Fixed_8sp[[i]][,6]))/var(Resu
lts_Fixed_8sp[[i]][,6])->R2_8sp[i]
}

R2_16sp<- vector(mode="numeric",length=Pool)   # R2 cv store (SR = 16)
for(i in 1:Pool){
1-((sum((Results_Fixed_16sp[[i]][,6] -
Results_Picante_16sp[[i]][,6])^2))/length(Results_Fixed_16sp[[i]][,6]))/var(Re
sults_Fixed_16sp[[i]][,6])->R2_16sp[i]
}
```

### Calculation of tree imbalance (Colless' index) #

```
require(apTreeshape)
Coll <- vector(mode="numeric",length=Pool)      # Colless' index store
for(i in 1:Pool){
colless(as.treeshape(Tree_List[[i]]))->Coll[i]
}
```

### Plotting results #

### Figure 1

```
require(vioplot)
vioplot(R2_2sp,R2_4sp,R2_8sp,R2_16sp, names=c("2","4","8","16"),col="khaki1")
title("Species pool n = 25")
```

### Figure 2

```
par(mfrow=c(1,4))

plot(1,type="n",ylim=c(-4,3),xlim=c(-4,3),xlab="ses.PD (ses.pd-fixed)",ylab="
ses.PD (Ses.pd-Picante)",main="Sample richness n = 2",axes=FALSE)
abline(0,1,col="grey")
axis(1,at=c(-10,-4,-3,-2,-1,0,1,2,3,10))
axis(2,at=c(-10,-4,-3,-2,-1,0,1,2,3,10))
axis(3,at=c(-100,10))
axis(4,at=c(-100,10))
```

```
for(i in 1:Pool){
points(Results_Fixed_2sp[[i]][,6],Results_Picante_2sp[[i]][,6],pch=19,cex=0.1)
}

plot(1,type="n",ylim=c(-4,3),xlim=c(-4,3),xlab="ses.PD (ses.pd-fixed)",ylab="
ses.PD (Ses.pd-Picante)",main="Sample richness n = 4",axes=FALSE)
abline(0,1,col="grey")
axis(1,at=c(-10,-4,-3,-2,-1,0,1,2,3,10))
axis(2,at=c(-10,-4,-3,-2,-1,0,1,2,3,10))
axis(3,at=c(-100,10))
axis(4,at=c(-100,10))
for(i in 1:Pool){
points(Results_Fixed_4sp[[i]][,6],Results_Picante_4sp[[i]][,6],pch=19,cex=0.1)
}

plot(1,type="n",ylim=c(-4,3),xlim=c(-4,3),xlab="ses.PD (ses.pd-fixed)",ylab="
ses.PD (Ses.pd-Picante)",main="Sample richness n = 8",axes=FALSE)
abline(0,1,col="grey")
axis(1,at=c(-10,-4,-3,-2,-1,0,1,2,3,10))
axis(2,at=c(-10,-4,-3,-2,-1,0,1,2,3,10))
axis(3,at=c(-100,10))
axis(4,at=c(-100,10))
for(i in 1:Pool){
points(Results_Fixed_8sp[[i]][,6],Results_Picante_8sp[[i]][,6],pch=19,cex=0.1)
}

plot(1,type="n",ylim=c(-4,3),xlim=c(-4,3),xlab="ses.PD (ses.pd-fixed)",ylab="
ses.PD (Ses.pd-Picante)",main="Sample richness n = 16",axes=FALSE)
abline(0,1,col="grey")
axis(1,at=c(-10,-4,-3,-2,-1,0,1,2,3,10))
axis(2,at=c(-10,-4,-3,-2,-1,0,1,2,3,10))
axis(3,at=c(-100,10))
axis(4,at=c(-100,10))
for(i in 1:Pool){
points(Results_Fixed_16sp[[i]][,6],Results_Picante_16sp[[i]][,6],pch=19,cex=0.
1)
}
```

### Figure 3

`(Coll-min(Coll))/(max(Coll)-min(Coll))->X1` # standardization of the Colless' index values between 0 (minimum value in the distribution) and 1 (maximum value in the distribution)

```
par(mfrow=c(1,4))
plot(R2_2sp~X1,pch=19,main="Sample richness n = 2",ylab="R2 cv",xlab="Colless'
index",xlim=c(0,1),ylim=c(min(c(R2_2sp,R2_4sp,R2_8sp,R2_16sp)),1))
plot(R2_4sp~X1,pch=19,main="Sample richness n = 4",ylab="R2 cv",xlab="Colless'
index",xlim=c(0,1),ylim=c(min(c(R2_2sp,R2_4sp,R2_8sp,R2_16sp)),1))
plot(R2_8sp~X1,pch=19,main="Sample richness n = 8",ylab="R2 cv",xlab="Colless'
index",xlim=c(0,1),ylim=c(min(c(R2_2sp,R2_4sp,R2_8sp,R2_16sp)),1))
plot(R2_16sp~X1,pch=19,main="Sample richness n = 16",ylab="R2 cv",xlab="Colless'
index",xlim=c(0,1),ylim=c(min(c(R2_2sp,R2_4sp,R2_8sp,R2_16sp)),1))
```

#### Supplementary figures referred in the main text of the paper

**Figure S1.** Violin plots showing the cross-validations R-squared scores for the comparisons between the ses.PD values derived from the functions *ses.pd*-Picante (Kembel et al., 2010) and its corrected version. Analyses were conducted using 500 simulated phylogenies per species pool (i.e. community data matrices).

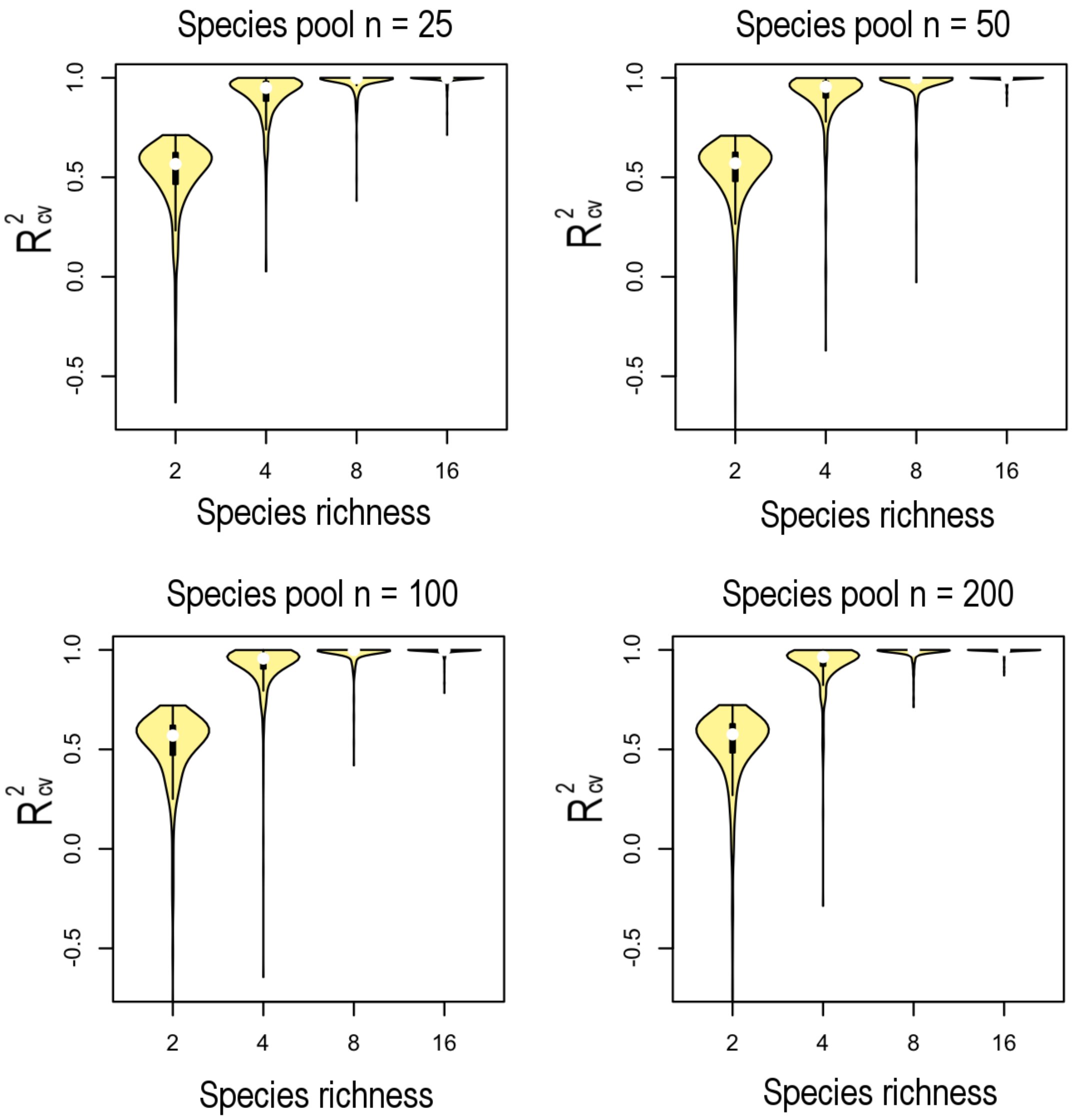

**Figure S2.** Scatter plots showing the relationship between ses.PD values (25,000 per plot) derived from the functions *ses.pd*-Picante (Kembel et al. 2010, y-axis) and its corrected version (x-axis). Analyses were conducted using datasets with species richness n = 2, 4, 8 and 16, respectively, species pools of (a) 25 species, (b) 50 species, (c) 100 species and (d) 200 species, and 500 simulated phylogenies per pool. The grey lines represent the expected 1:1 relationship.

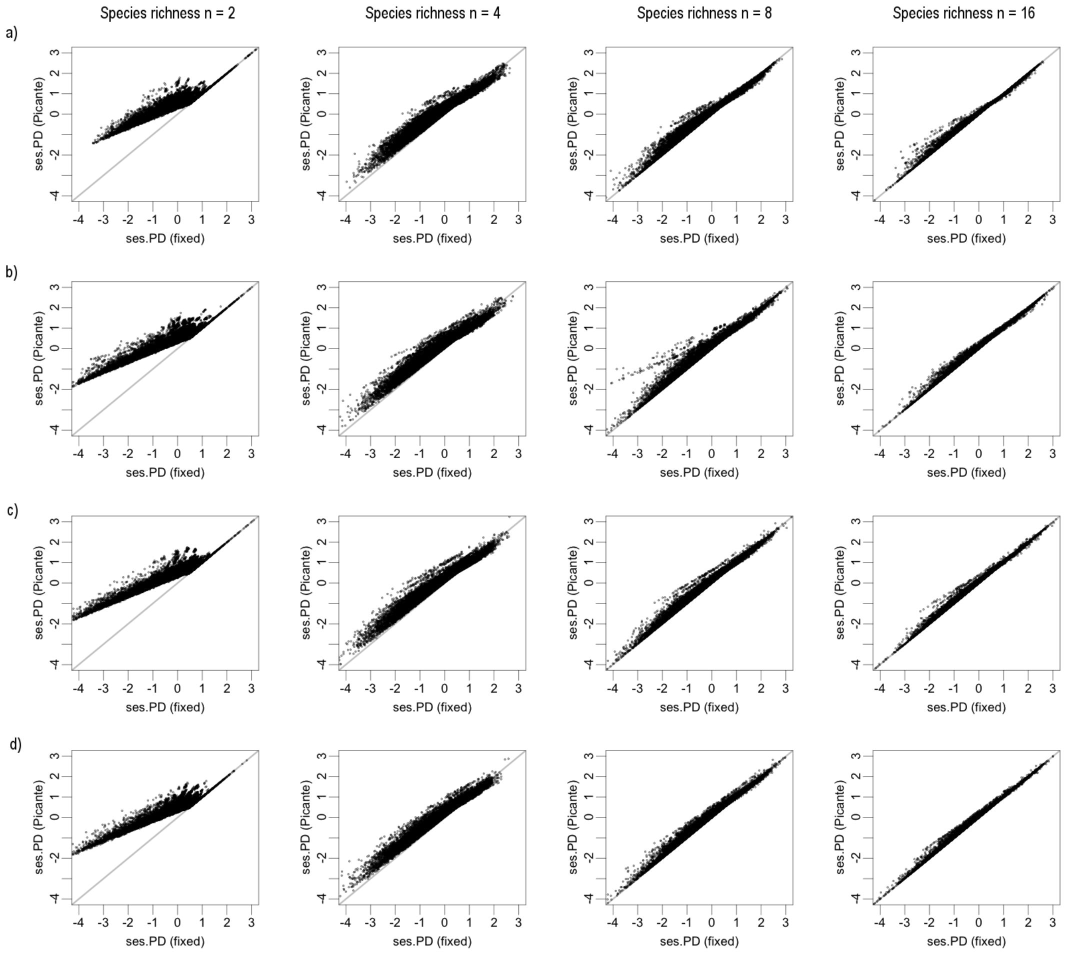

**Figure S3.** Relationship between cross-validation R-squared scores (y-axis) and tree imbalance (i.e. Colless' index, x-axis). Analyses were conducted using datasets with species = 2, 4, 8 and 16, respectively, species pools of (a) 25 species, (b) 50 species, (c) 100 species and (d) 200 species and 500 simulated phylogenies per pool. The higher the value of the Colless' index, the higher the imbalance of the phylogeny (values scaled between 0 and 1).

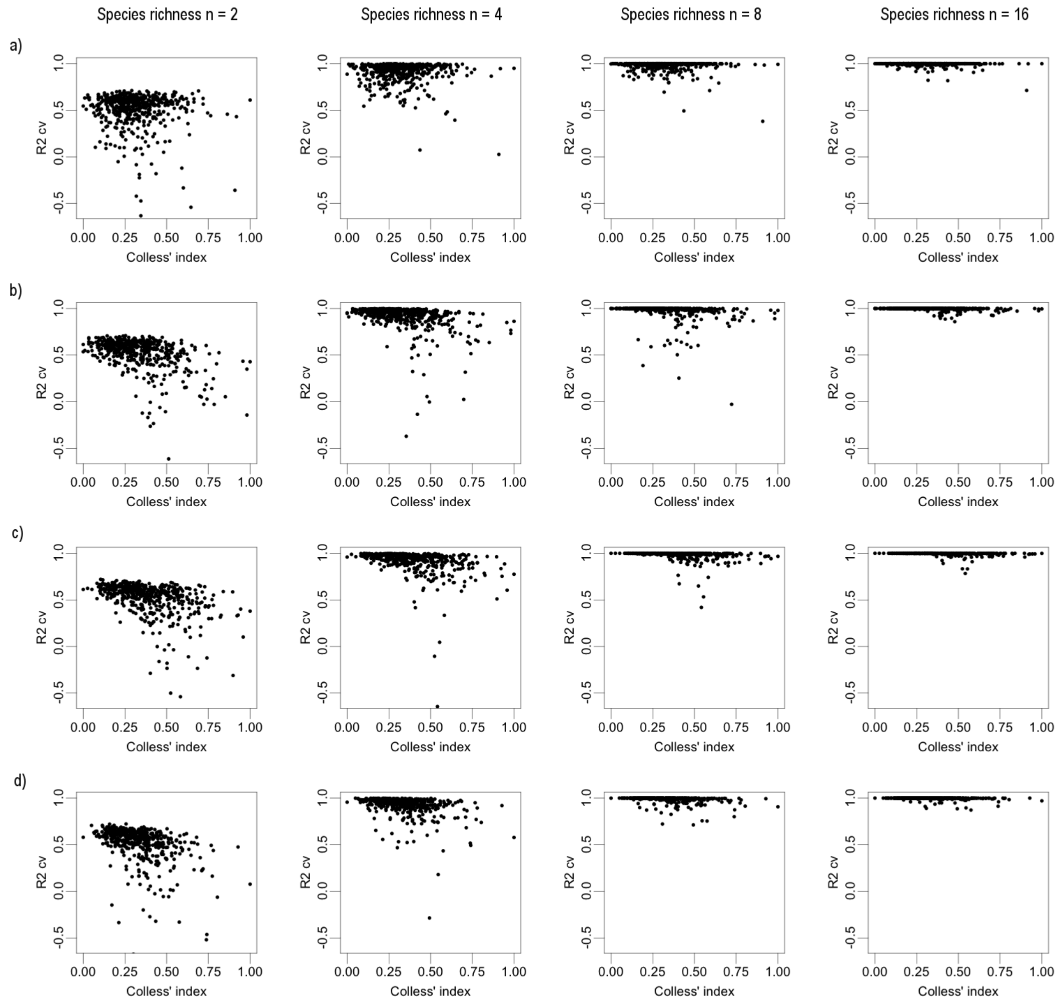

#### <u>Corrected version of the *ses.pd*-Picante function</u>

```
ses.pd.fixed<-function(samp, tree, null.model = c("taxa.labels", "richness",
"frequency", "sample.pool", "phylogeny.pool", "independentswap", "trialswap"),
runs = 999, iterations = 1000, include.root = TRUE)
{

if(include.root == TRUE) {
pd.obs <- as.vector(pd(samp, tree, include.root = TRUE)$PD)
null.model <- match.arg(null.model)

pd.rand <- switch(null.model, taxa.labels = t(replicate(runs,
as.vector(pd(samp, tipShuffle(tree), include.root = TRUE)$PD))), richness =
t(replicate(runs,as.vector(pd(randomizeMatrix(samp, null.model = "richness"),
tree, include.root = TRUE)$PD))), frequency = t(replicate(runs, as.vector
(pd(randomizeMatrix(samp,null.model = "frequency"), tree, include.root =
TRUE)$PD))), sample.pool = t(replicate(runs,as.vector(pd(randomizeMatrix(samp,
null.model = "richness"), tree, include.root = TRUE)$PD))), phylogeny.pool =
t(replicate(runs, as.vector(pd(randomizeMatrix(samp, null.model =
"richness"),tipShuffle(tree), include.root = TRUE)$PD))), independentswap =
t(replicate(runs,as.vector(pd(randomizeMatrix(samp, null.model =
"independentswap", iterations), tree, include.root = TRUE)$PD))), trialswap =
t(replicate(runs,as.vector(pd(randomizeMatrix(samp, null.model = "trialswap",
iterations), tree, include.root = TRUE)$PD))))

pd.rand.mean <- apply(X = pd.rand, MARGIN = 2, FUN = mean, na.rm = TRUE)
pd.rand.sd <- apply(X = pd.rand, MARGIN = 2, FUN = sd, na.rm = TRUE)
pd.obs.z <- (pd.obs - pd.rand.mean)/pd.rand.sd
pd.obs.rank <- apply(X = rbind(pd.obs, pd.rand), MARGIN = 2, FUN = rank)[1, ]
pd.obs.rank <- ifelse(is.na(pd.rand.mean), NA, pd.obs.rank)
Result<-data.frame(ntaxa = specnumber(samp), pd.obs, pd.rand.mean,
pd.rand.sd, pd.obs.rank, pd.obs.z, pd.obs.p = pd.obs.rank/(runs + 1), runs =
runs, row.names = row.names(samp))
return(Result)
}

if(include.root == FALSE) {
pd.obs <- as.vector(pd(samp, tree, include.root = FALSE)$PD)
null.model <- match.arg(null.model)

pd.rand <- switch(null.model, taxa.labels = t(replicate(runs,
as.vector(pd(samp, tipShuffle(tree), include.root = FALSE)$PD))), richness =
t(replicate(runs,as.vector(pd(randomizeMatrix(samp, null.model = "richness"),
tree, include.root = FALSE)$PD))), frequency = t(replicate(runs, as.vector
```

```
(pd(randomizeMatrix(samp,null.model = "frequency"), tree, include.root =
FALSE)$PD))),sample.pool = t(replicate(runs,as.vector(pd(randomizeMatrix(samp,
null.model = "richness"), tree, include.root = FALSE)$PD))), phylogeny.pool =
t(replicate(runs, as.vector(pd(randomizeMatrix(samp, null.model =
"richness"),tipShuffle(tree), include.root = FALSE)$PD))), independentswap =
t(replicate(runs,as.vector(pd(randomizeMatrix(samp, null.model =
"independentswap", iterations), tree, include.root = FALSE)$PD))), trialswap =
t(replicate(runs,as.vector(pd(randomizeMatrix(samp, null.model = "trialswap",
iterations), tree, include.root = FALSE)$PD))))

pd.rand.mean <- apply(X = pd.rand, MARGIN = 2, FUN = mean, na.rm = FALSE)
pd.rand.sd <- apply(X = pd.rand, MARGIN = 2, FUN = sd, na.rm = FALSE)
pd.obs.z <- (pd.obs - pd.rand.mean)/pd.rand.sd
pd.obs.rank <- apply(X = rbind(pd.obs, pd.rand), MARGIN = 2, FUN = rank)[1, ]
pd.obs.rank <- ifelse(is.na(pd.rand.mean), NA, pd.obs.rank)
Result<-data.frame(ntaxa = specnumber(samp), pd.obs, pd.rand.mean,
pd.rand.sd, pd.obs.rank, pd.obs.z, pd.obs.p = pd.obs.rank/(runs + 1), runs =
runs, row.names = row.names(samp))
return(Result)
}
}
```
